# Assessing Conformation Validity and Rationality of Deep Learning-Generated 3D Molecules

**DOI:** 10.1101/2024.11.10.622844

**Authors:** Fan Fan, Bin Xi, Xianghu Meng, Han Wang, Bowen Zhang, Qingbo Xu, Wei Feng, Xiaoman Wang, Hongbo Zhang, Feng Zhou, Zhenming Liu, Wenbiao Zhou, Bo Huang

**Affiliations:** Beijing StoneWise Technology Co Ltd., Haidian Street #15, Beijing, 100080, China; State Key Laboratory of Natural and Biomimetic Drugs, School of Pharmaceutical Sciences, Peking University, Beijing, China

**Author notes:** These authors contributed equally to this work (Fan Fan, Bin Xi, Xianghu Meng).

## Abstract

Recent advancements in artificial intelligence (AI) have revolutionized the field of 3D molecule generation. However, the lack of effective evaluation methods for 3D conformations limits further improvements. Current techniques, in order to achieve the necessary speed for evaluating large numbers of AI-generated molecules, often rely on empirical geometric metrics that do not adequately capture various conformational anomalies, or on molecular mechanics (MM) energy metrics that exhibit low accuracy and lack atomic or torsional details. To address this gap, we propose a two-stage approach that achieves both high speed and quantum mechanical (QM) level accuracy. The first stage, termed the validity test, employs an AI-derived force field to identify atoms with elevated energy resulting from implausible neighboring environments. The second stage, known as the rationality test, utilizes a deep learning network trained on data with density functional theory (DFT) accuracy to detect rotatable bonds with high torsional energies. To demonstrate the functionality of our evaluation system, we applied our approach to five recently reported 3D molecule generation AI models across 102 targets in Directory of Useful Decoys-Enhanced (DUD-E) dataset. To facilitate accessibility for the academic community, our method is available as an open-source package.

## 1. Introduction

Recent advances in deep learning generative models have begun to demonstrate the ability to generate small molecules with 3D conformations based on a given protein pocket ^1^. Both auto-regressive models and diffusion models have shown this capability^2-7^. However, these AI models continue facing challenges of generating physically implausible 3D conformations. A number of studies have reported the production of abnormal conformations, characterized by steric clashes, twisted structures, and the misplacement of hydrogen atoms^3,8^.

To quantify the issues of abnormal conformation and thereby guide AI model training, two types of evaluation methods are frequently adopted: (1) geometry-based and (2) energy-based conformation assessment. Geometric approaches typically examine bond lengths, bond angles, the overlap of Van der Waals radii of two atoms^9,10^, or the redocking root-mean-squared deviation (RMSD)^8^. However, these geometric metrics are intrinsically limited by the lack of an energy criterion, which can lead to misleading results. For instance, two conformations with a small RMSD may have substantially different energies. Additionally, geometric approaches may use Kullback-Leibler or Jensen-Shannon divergence to evaluate AI-generated conformations against reference molecules by analyzing geometric features like bond lengths and angles^3,6^. The reliability of such measurement heavily relies on the reference set, whose limited coverage may cause misleading when evaluating molecules which are not similar to those in the reference set. Energy-based metrics often strive to achieve a balance between accuracy and time efficiency. High-precision calculations, such as density functional theory (DFT)^11^ techniques, can be quite time-consuming. On the contrary, time-efficient alternatives such as semi-empirical^12^ and molecular mechanics (MM) methods^13-16^ often have relatively low accuracy. More importantly, most current energy-based methods provide only global energies at the molecular level, lacking detailed atom-level or torsion-level energy assessments. This limitation presents a significant challenge for AI algorithm designers, who require detailed insights into the positioning of individual atoms.

In addition to the challenges related to evaluation tactics, the debate over evaluation strategies for AI-generated conformations further complicate the situation. Some studies advocate for pre-refinement evaluation, which involves the direct assessment of AI-generated conformations without MM refinement^2,4,8^, while others propose post-refinement evaluation, which involves a preliminary MM optimization with the protein pocket fixed prior to assessment^3,7^. Both of the pre-refinement and post-refinement evaluations have their pros and cons. Pre-refinement evaluations can reveal model deficiencies that may be concealed by force field optimization; however, they often lack the sensitivity needed for detailed analysis, especially when quantum mechanics (QM) computations are involved. For example, DFT assessments of torsion energies can be skewed by abnormal bond lengths. Additionally, a model’s performance in pre-refinement evaluations doesn’t necessarily reflect its overall utility, as medicinal chemists might prefer a model that, despite initial shortcomings, provides more rational conformations after cost-effective force field-based refinement. This emphasizes the complexity of model evaluation in practical applications.

Given the pros and cons of these evaluation strategies and the limitations of existing tactics, there is an urgent need to develop a systematic framework for evaluating AI generated conformations.

To overcome the limitations in current evaluation methodologies, we propose a two-stage procedure consisting of validity and rationality tests for AI generated conformations before and after force-field refinement, respectively. For pre-refinement conformations, we determine whether they are valid by detecting abnormal conformations at atomic level with energy-based metrics. The tool developed for this purpose, named HEAD (<u>h</u>igh-<u>e</u>nergy <u>a</u>tom <u>d</u>etection), utilizes the network of machine learning force fields (MLFFs) to compute the atomic energies corresponding to each atom’s local environment. HEAD shows higher efficiency in detecting abnormal conformations compared to the widely used benchmark method PoseBusters^8^, which mainly rely on geometry-based metrics. However, a force-field valid conformation may still be irrational because of high torsion energy. To this end, we propose a rationality test designed to quantify the disparity between post-refinement conformations and low-torsional energy conformations. The tool developed for this purpose, named TED (<u>t</u>orsional <u>e</u>nergy <u>d</u>escriptor), is mainly composed by a deep learning-based torsion energy prediction model (referred to as the TED-Model henceforth). To train TED-Model, we developed two data sets including a pretraining set containing semi-empirical torsion energy data for six million torsion fragments and a training set containing double-hybrid DFT data for 100,000 torsion fragments. TED demonstrates superior accuracy compared to GFN2-xTB when evaluated on a data set of 5,000 torsional fragments without information leakage. To illustrate the application of our evaluation system, we evaluated five recently reported 3D molecule generative models including Lingo3DMolv2^3^, Pocket2Mol^5^, PocketFlow^2^, TargetDiff^6^, and PMDM^4^. Each model generated around 1,000 molecules per target across 102 targets from Directory of Useful Decoys-Enhanced (DUD-E) dataset^17^ and underwent both validity and rationality tests. To facilitate the use of our evaluation system, we have made the HEAD and TED models accessible via link https://github.com/stonewiseAIDrugDesign/HEAD_TED.

## 2. Results and Discussions

In this section, we first describe the construction of our HEAD and TED modules (Figure 1), outlining the fundamental logic behind their design and presenting testing results that demonstrate their reliability. We then report an evaluation test in which HEAD and TED were applied to five recently reported 3D molecule generation models: Lingo3DMolv2, TargetDiff, Pocket2Mol, PocketFlow, and PMDM.

**Figure 1.**
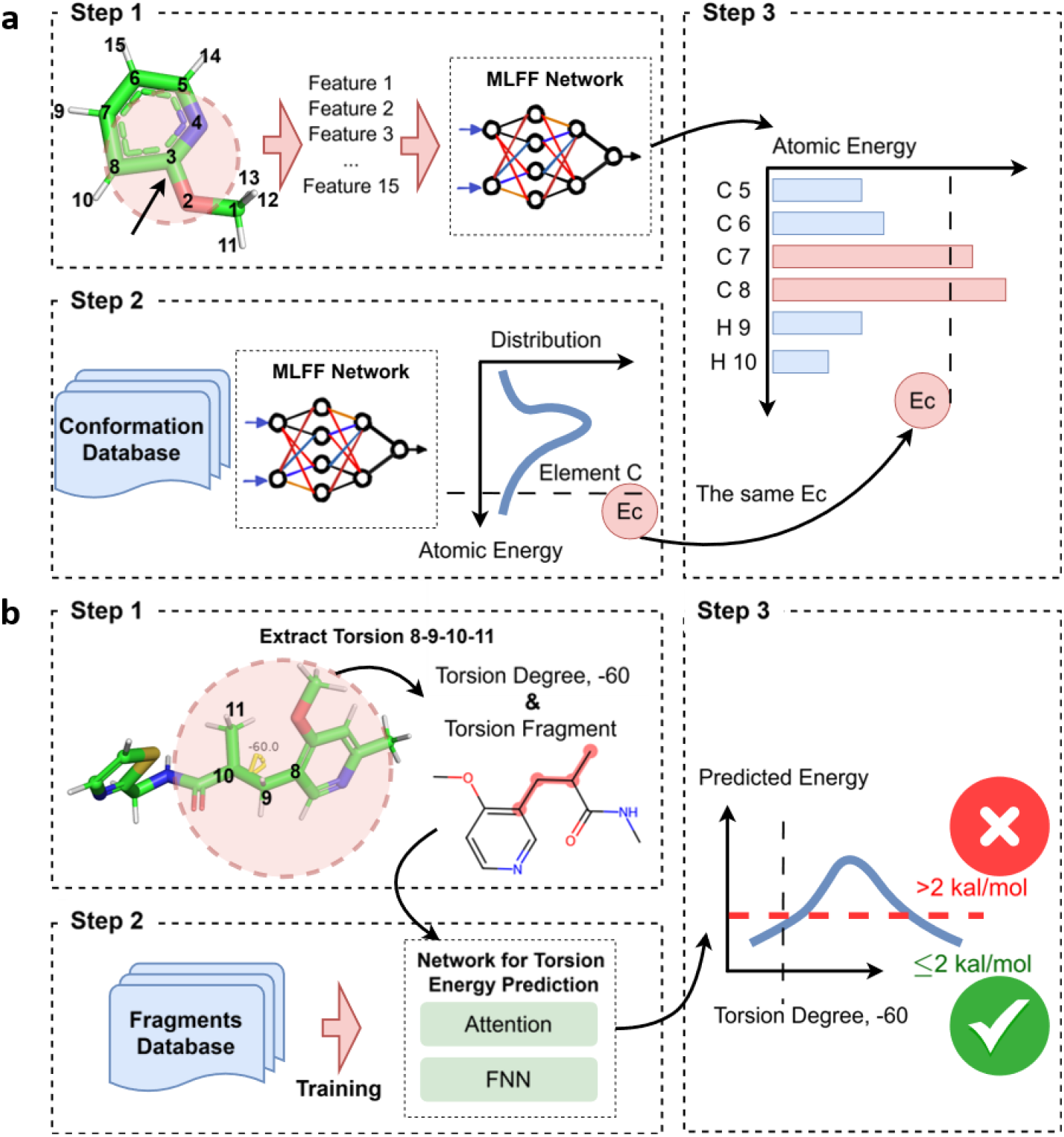
Schematic diagram of the HEAD and TED modules for 3D conformation evaluation. (a) Overview of the HEAD module. It assesses conformation validity by calculating atomic energies (step 1), establishing thresholds for anomaly detection (step 2), and classifying conformations based on energy comparisons against the threshold (step 3). The threshold is represented as *E*_*c*_ within a pink circle. (b) Overview of the TED module. It evaluates conformation rationality by decomposing the structure into torsion fragments (step 1), predicting torsion energies for the target rotatable bond within each fragment (step 2), and assigning a binary label based on an energy threshold of 2 kcal/mol (step 3).

### 2.1 Development of HEAD

The HEAD module is designed to assess the quality of AI generated 3D molecular conformations without MM optimization. It directly takes an AI-generated conformation as input and quantitatively identifies atoms responsible for the anomalies of the conformation. This module is developed through three steps, as illustrated in Figure 1a. The first step involves computing atomic energies using MLFFs. The MLFFs used in this study is ANI-2x^18,19^. The second step focuses on establishing thresholds to classify energy values that are significantly higher than normal. This is accomplished by applying the MLFF method to molecules from classical databases to obtain energy distributions for each elemental type. Based on these distributions, thresholds for each element are established. The third step involves comparing the atomic energies in a given molecule with the thresholds of the corresponding elements, resulting in a binary label indicating whether the molecular conformation is abnormal. Detailed information regarding the three steps is provided in the Methods section 4.1.

To evaluate the accuracy of the HEAD approach, we utilized three distinct databases. First, LigBoundConf dataset^20^, was used as a benchmark for valid conformations of ligand binding with a protein. This dataset includes drug-like molecules that are sourced from the Protein Data Bank (PDB)^21^ and refined with force field in the presence of binding proteins. Second, the Cambridge Structural Dataset (CSD)^22^, was used as a benchmark for valid conformations of ligand without protein binding. This dataset is comprised of high-quality experimentally determined small molecule crystal structures. Both the CSD and LigBoundConf acted as positive controls, enabling us to determine whether HEAD incorrectly classifies valid conformations as invalid.

Additionally, we introduced the third database, GM-5K, which consists of randomly selected AI-generated molecules provided by Lingo3DMolv2, TargetDiff, Pocket2Mol, PocketFlow, and PMDM. Because GM-5K dataset contains conformations of varying quality, we employed some calculations to label the conformation quality. Specifically, we employed MMFF94 force field^14^ for ligand geometry optimization with protein pocket fixed and conducted QM-level (revDSD-PBEP86-D3(BJ)/def2-TZVPP)^23-25^ single point energy computations for the conformations before and after optimization. The energy difference between original and optimized conformations, i.e., 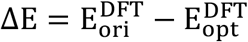, was used as quality indicator of the original conformation before optimization. GM-5K dataset supported the test of HEAD’s ability to distinguish abnormal conformations from valid ones.

Due to the presence of elements not supported by ANI-2x in some molecules from the three aforementioned datasets, we applied a screening criterion that limits inclusion to molecules composed of commonly occurring elements: hydrogen (H), carbon (C), nitrogen (N), oxygen (O), fluorine (F), sulfur (S), and chlorine (Cl). After applying this criterion, we retained 159,948 molecules from the CSD, 6,209 from LigBoundConf, and 5,000 from GM-5K.

Regarding benchmark technology, we compared HEAD with PoseBusters^8^. PoseBusters is a recently released test suite enabling implausible conformation identification. It utilizes the RDKit^26^ toolset to evaluate input molecular conformations by analyzing geometry metrics such as bond lengths, bond angles, aromatic ring planarity, double bond planarity, and internal steric clashes. It also employs an energy metric that determines whether the input molecule’s MM-level energy exceeds 100 times the average energy of 50 force field-optimized conformations. We assessed the performance of HEAD and PoseBusters from three perspectives: (1) recall rate in positive control datasets, (2) F1 score in the GM-5K dataset, and (3) speed.

We first tested the recall rate of HEAD and PoseBusters on CSD and LigBoundConf, which indicates how much the approaches intend to incorrectly classifies valid conformations as invalid. As shown in Figure 2a, the valid ratio for HEAD and PoseBusters on both LigBoundConf and CSD dataset are all over 95%. It is noteworthy that, although HEAD and PoseBusters show similar performance, HEAD is around 30 times faster than PoseBusters. This makes HEAD approach more suitable for high-throughput evaluation tasks. A detailed speed comparison can be found in Extended Data Table 1.

**Figure 2.**
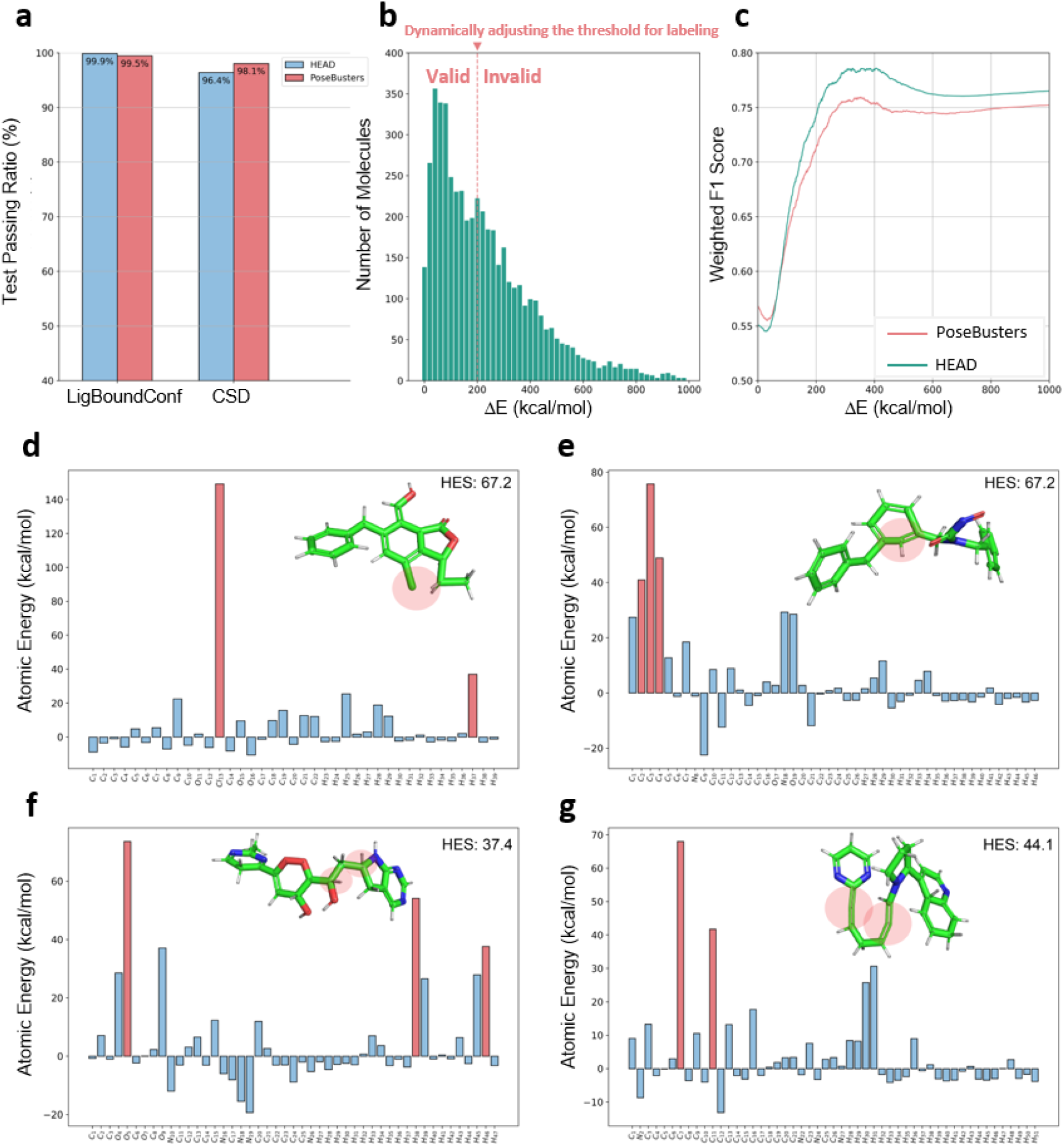
Comparison of HEAD and PoseBusters. (a) Valid conformation recall rates of HEAD and PoseBusters tested using CSD and LigBoundConf. Molecular conformations in CSD and LigBoundConf are all considered as valid, representing valid conformations without and with protein binding, respectively. (b) Histogram distribution of molecules in GM-5K dataset across ΔE with the bin size of 20 kcal/mol. 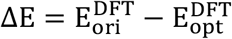.The red dashed line indicates the threshold value used to classify molecular conformations as valid or invalid. Molecules with ΔE values below the threshold are labeled as valid, while those above the threshold are labeled as invalid. The dynamic adjustment of the threshold value leads to corresponding changes in valid and invalid labels. (c) Weighted F1 scores of the HEAD and PoseBusters across varying labeling thresholds based on ΔE in the GM-5K dataset. The threshold was systematically varied from 0 to 1000 kcal/mol in increments of 1 kcal/mol, resulting in a series of weighted F1 scores. The weighted F1 score is computed by first evaluating the F1 scores for both valid and invalid classes. Subsequently, these F1 scores are averaged considering the number of samples in each class as weights. Panel (d) to (g) showcase four representative abnormal geometries with their atomic energies, including steric clash (d), twisted ring structure (e), misplacement of hydrogen atom (f), and valence violation (g). The atom names are listed on the horizontal axis and their energy values are on the vertical axis. The detected high atomic energies (in red bars) consistently reflect the invalid region (circled in red) of each conformation.

Next, we checked the weighted F1 scores of the HEAD and PoseBusters models on the GM-5K dataset to evaluate their ability to discriminate between valid and invalid 3D molecular conformations. GM-5K contains molecules with ΔE varying from 0 kcal/mol to 1000 kcal/mol, shown as the histogram distribution displayed in Figure 2b. To compare the discrimination powers of HEAD and PoseBusters, we need to assign a valid-invalid binary label to every molecule in GM-5K based on its ΔE value. Specifically, molecules with ΔE values over a specified threshold should be classified as invalid, while those with values below the threshold should be classified as valid. Since there is no universal threshold for classifying conformation validity, we systematically varied the threshold from 0 to 1000 in increments of 1 kcal/mol. As we adjusted the labeling threshold, as illustrated in Figure 2b, the corresponding valid and invalid labels were updated accordingly. This process resulted in a set of F1 scores that correspond to the variations of labeling thresholds. The results, plotted in Figure 2c, shows that HEAD notably outperforms PoseBusters in identifying abnormal conformations with ΔE values between 200 kcal/mol and 600 kcal/mol. For the conformations with ΔE values outside this region, the two methods exhibited comparable performance. Some example cases that can be detected by HEAD but not PoseBusters are shown in Figure 2d-g. The examples include abnormal bond lengths, bond angles, steric clashes, valence violation, misplacement of hydrogens, twisted rings and etc. The reason HEAD outperforms PoseBusters in detecting anomalous conformations may be related to the missing definitions of geometric anomalies in PoseBusters. Specifically, anomaly detection is more effective in energy space than in real space. It is highly difficult to enumerate all geometric anomalies in real space. In contrast, all abnormal situations correspond to high energy responses.

It is also notable that for conformations with relatively small anomalies, indicated by low ΔE values, both PoseBusters and HEAD experience a sharp drop in F1-score. This trend indicates the limitations of these methods for detecting anomalies within this range. The potential causes of these limitations will be discussed in the Discussion section.

### 2.2 Development of TED

The TED module is developed to evaluate the quality of AI generated 3D molecular conformations after MM based optimization. It accepts a MM-refined conformation as input, quantitatively predicts the torsion energies for each rotatable bond that excludes hydrogens, and subsequently outputs a binary label indicating whether the input 3D conformation is abnormal. This module works in three steps, as illustrated in Figure 1b. The first step involves decomposing a given conformation into a series of torsion fragments, each containing a target rotatable bond and its necessary local chemical environment. The second step predicts the torsion energy curve for the target rotatable bond in the torsion fragment using a deep learning model (i.e. TED-Model) that employs an attention mechanism. TED-Model was trained with 6 million torsion fragments with semi-empirical level energy data and fine-tuned with 100,000 torsion fragments with DFT level energy data. The third step assigns a rational-irrational binary label to the conformation by checking whether any of its rotatable bonds have a torsion energy exceeding 2 kcal/mol, a threshold employed in a previous study^27^. Comprehensive details regarding the development of the TED module, including TED-Model training and dataset construction, are provided in the Method section 4.2.

To evaluate the performance of our TED-Model, we compared its predictions with those obtained using the semiempirical GFN2-xTB method^12^. This assessment utilized the DFT-5K dataset, which contains 5,000 unique torsion fragments which are not included in the training set of our model for the mitigation of information leakage. Each torsion fragment in DFT-5K has 24 conformers generated and labeled with DFT-level energies using approaches described in Methods section 4.2.1.

To assess the alignment of our TED-Model’s predictions with DFT values relative to GFN2-xTB, we computed Pearson correlations on a per-torsion-fragment basis using the DFT-5K dataset. For each torsion fragment, we analyzed the correlations between the energies predicted by our model across 24 conformers and the corresponding DFT energies, as well as the correlations between GFN2-xTB predictions and DFT values. The distribution of these Pearson correlations is presented in Figure 3a. Our model exhibits a stronger agreement with DFT values, with a per-torsion-fragment average Pearson correlation of 0.84, compared to 0.63 for GFN2-xTB. This superior performance can be attributed to our model’s finetuning using DFT data, which enables it to effectively address scenarios where GFN2-xTB encounters challenges. Specifically, GFN2-xTB exhibits limitations in accurately characterizing anisotropy in electrostatic potential and polarization effects. For instance, the torsion fragment shown in Figure 3b demonstrates that GFN2-xTB overestimates the torsion energy at ±180°, primarily due to its inadequate treatment of sigma-hole interactions between sulfur and the oxygen of the carbonyl group. Additionally, Figure 3c demonstrates GFN2-xTB’s tendency to overestimate lone-pair repulsion, leading to an increase in torsion energy at ±180°.

**Figure 3.**
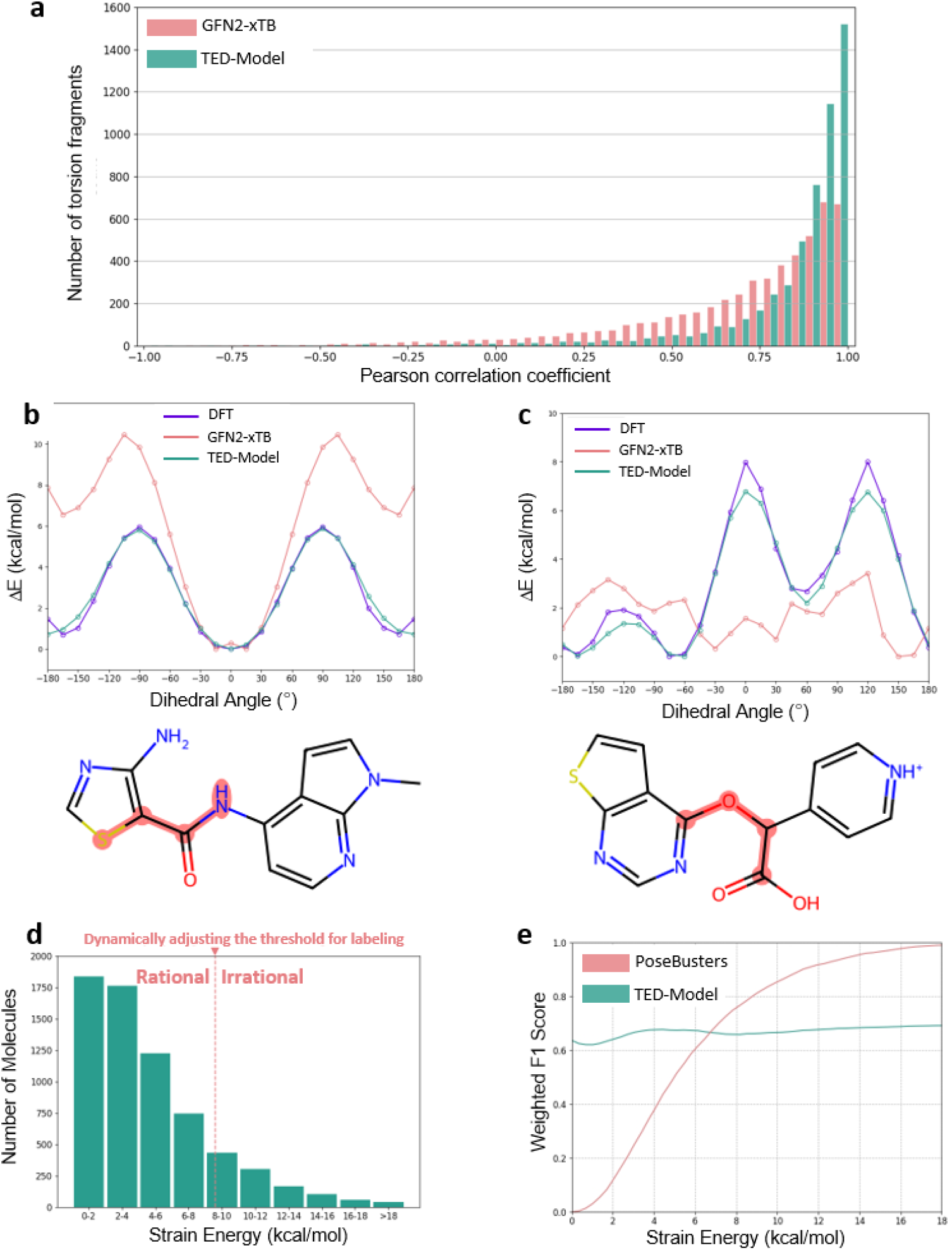
Evaluation of TED in torsion energy predictions and irrational conformation discrimination. (a) Histogram distribution of torsion fragments in the DFT-5K dataset across Pearson correlation coefficients between DFT values and predictions from our TED-Model and GFN2-xTB. Torsion energies were computed for the 24 conformations of each torsion fragment in DFT-5K dataset. Pearson correlation coefficients between both methods and DFT were calculated on a per-torsion-fragment basis respectively. (b) Case illustration of our model’s superior performance to GFN2-xTB’s in the scenario of sigma-hole interactions. There is a sigma-hole associated interaction between the sulfur of thiazole group and the oxygen of the carbonyl group when dihedral angle of S-C-C-N is ±180°. (c) Case illustration of our model’s superior performance to GFN2-xTB’s in the scenario of lone-pair repulsion. There is a lone-pair repulsion between the nitrogen of the thienopyridine group and the oxygen of the carboxy group when dihedral angle of C-O-C-C is ±180°. For panels (b) and (c), the atoms defining the dihedral angle are marked with red circles. (d) Histogram distribution of molecules in LigBoundConf dataset across strain energy. The red dashed line indicates the threshold value used to classify molecular conformations as rational or irrational. Molecules with strain energy above the threshold are labeled as irrational, while those below the threshold are labeled as rational. The dynamic adjustment of the threshold value leads to corresponding changes in rational and irrational labels. (e) Comparison of weighted F1 scores between our TED method and PoseBusters on the LigBoundConf dataset, with varying thresholds for rational and irrational labeling. The threshold was systematically varied from 0 to 18 kcal/mol in increments of 0.2 kcal/mol, resulting in a series of weighted F1 scores. The weighted F1 score is computed by calculating the F1 scores for both the rational and irrational classes and then averaging these scores using the number of samples in each class as weights.

Next, we compared our TED method with PoseBusters^8^, both serving as discriminators in distinguishing irrational conformations arising from ligand conformational strain energy issues. To facilitate this comparison, we employed the LigBoundConf dataset^20^, which contains 8,145 small molecules with strain energy provided as the energy difference between bound conformations in proteins and global minimum energy conformations without protein binding.

We assessed the weighted F1 scores of both TED and PoseBusters on the LigBoundConf dataset to evaluate their discriminative power. The weighted F1 scores were calculated using varying thresholds for rational and irrational labeling (Figure 3d), consistent with our methodology for testing HEAD and PoseBusters on GM-5K, as described in section 2.1.

The results are illustrated in Figure 3e. PoseBusters exhibits a weighted F1 score of one at elevated strain energy thresholds, but this score sharply declines to zero as the strain energy threshold decreases. This trend indicates that PoseBusters has no discriminative power. Further examination reveals that all conformations in the LigBoundConf dataset passed the PoseBusters test. The underlying reason for this is that the LigBoundConf conformations underwent force field optimization, which minimizes the geometric issues that PoseBusters aims to detect.

In contrast, TED demonstrates a relatively stable discriminative power, achieving weighted F1 scores above 0.6. It is important to note that strain energy differs from torsion energy, and this discrepancy may contribute to the observed mild F1 scores. We will discuss this later in the Discussion section.

### 2.3 Evaluation of Molecule Generative Models

In this section, we demonstrated the application of the HEAD&TED system in evaluating AI-generated molecules. Five recently reported AI generative models were included in this assessment. Specifically, Lingo3DMolv2, Pocket2Mol, and PocketFlow were selected to represent autoregressive models, while PMDM and TargetDiff were chosen as representatives of diffusion models. The evaluation was conducted on 102 targets from the DUD-E dataset, with each model configured to generate 1,000 unique molecules per target. Some models were unable to generate 1,000 unique molecules for some of the 102 targets with reasonable resource consumption; further details can be found in the Supplementary Information Part 2 and 3. All the AI-generated molecules were submitted to HEAD and PoseBusters for conformation validity test. They were then refined using force field OPLS3e^28^ with protein pocket fixed and then submitted to TED for conformation rationality test.

Prior to evaluating the conformational quality of the generated molecules, we need to emphasize the necessity of eliminating those exhibiting low level of drug-likeness or poor synthetic accessibility. This approach is supported by below observations. Even though the molecules displayed in Extended Data Figure 1 successfully met the criteria set by PoseBusters, HEAD, and TED tests, they were not considered drug candidates due to their low level of drug-likeness. This was reflected in their low Quantitative Estimate of Drug-likeness (QED) scores^29^ and poor synthetic accessibility, as indicated by high Synthetic Accessibility Scores (SAS)^30^. Furthermore, we examined the distribution of HEAD and TED passing rates against QED and SAS for molecules produced by the AI models under test (Extended Data Figure 2). To establish the drug-like region, we defined criteria of a QED score of 0.3 or higher and a SAS score of 5 or lower, which collectively account for over 75% of the molecules listed in DrugBank^31^. As shown in Extended Data Figure 2, all the AI models under test have generated molecules which are not drug-like but show high passing rate of HEAD or TED. This observation further emphasizes the necessity of eliminating molecules outside drug-like region before conformation quality evaluation. Otherwise, the evaluation results would be contaminated.

Building on the elimination of molecules with QED lower than 0.3 and SAS higher than 5, we conducted conformation quality evaluation using both PoseBusters and our HEAD&TED. The conformation quality was analyzed on a per-target basis and defined as the percentage of molecules passing the specific test (Figure 4). The histogram distribution of target counts across PoseBusters passing rates for each AI model is shown in Figure 4a. Notably, Lingo3DMolv2 demonstrated superior performance, showing a distribution skewed towards high passing rate with a mean value of 76.4%. In contrast, other models, including Pocket2Mol, PocketFlow, TargetDiff, and PMDM, exhibited mean PoseBusters passing rates of 66.8%, 50.6%, 64.6%, and 51.6%, respectively. The HEAD validity test produced results comparable to those of PoseBusters (Figure 4b), while the TED test indicated that although PocketFlow scored poorly in validity assessments, it excelled in rationality testing compared to other models (Figure 4c). To further elucidate these findings, we examined the property distribution of molecules generated by different models (Extended Data Figure 3). The favorable performance of PocketFlow in the TED rationality test may be attributed to its generation of molecules having fewer rotatable bonds, lower molecular weights, and fewer chiral centers compared to other models.

**Figure 4.**
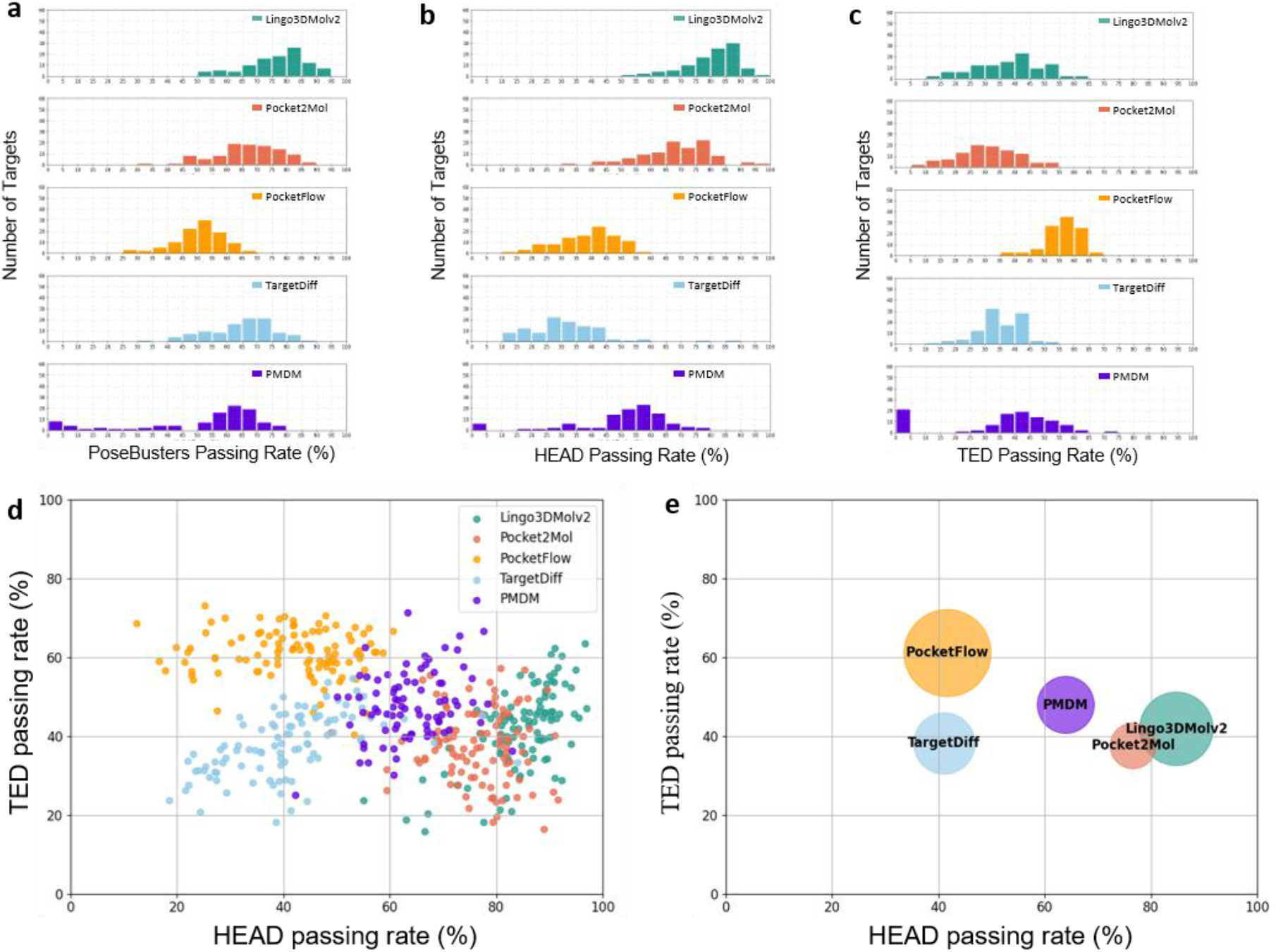
Evaluation of conformation quality for AI generated models. (a) – (c) Distributions of target counts based on the passing rates of PoseBusters, HEAD, and TED, with histograms presented in panels a, b, and c, respectively. (d) Scatter plot of HEAD and TED passing rates on a per-target basis, with points colored according to the AI models. (e) Bubble chart providing a comprehensive evaluation of AI models, where the position of each bubble centroid corresponds to the per-target average of HEAD and TED passing rates, and the size of the bubble reflects the percentage of drug-like molecules, defined as those with a QED greater than 0.3 and an SAS below 5.

The HEAD powered validity test reveals the limitations of the AI model which causing the gap between AI generated conformations and those refined by force field. However, due to the low cost and high speed of force field-based optimization, this gap can be mitigated by performing such optimization for all AI-generated molecules with pocket fixed. The critical question then becomes whether the force field-refined conformations truly align with the low-torsion energy criteria that drug designers consider acceptable, which is reflected in the TED powered rationality test. To compile a comprehensive view of these AI molecule generation models, we plotted their HEAD passing rates and TED passing rates on a per-target basis (Figure 4d). We then considered points from the same AI model as a cluster and used the centroid of each cluster to represent it. These centroids were then transformed into bubbles, with their sizes reflecting the percentage of drug-like molecules, specifically those with a QED greater than 0.3 and an SAS below 5. A larger bubble positioned in the upper right corner of Figure 4d indicates better overall performance of the model. As observed, no single model outperformed all others comprehensively. However, PocketFlow and Lingo3DMolv2 have the potential to achieve superior performance by improving their validity and rationality, respectively.

## 3. Discussion

In this work, we divided the evaluation of AI-generated 3D molecular conformations into two stages: validity and rationality tests. The validity test measures how closely AI-generated conformations align with force field-refined conformations, while the rationality test assesses how closely a force field-refined AI-generated conformation approaches a low-torsional energy conformation. To support these evaluations, we introduced two tools: HEAD, powered by an AI-driven force field, and TED, which utilizes an AI torsion energy prediction network. Unlike traditional methods that primarily rely on geometric measures or molecular-level energy metrics, HEAD provides enhanced granularity through atomic-level energy metrics, and TED improves interpretability by identifying rotatable bonds with high torsion energy. Given the critical role that evaluation frameworks play in guiding the iterative development of AI models, we hope that our methodology can facilitate the continuous improvement of AI-based 3D molecular generation models, steering them toward the generation of more valid and rational conformations.

Regarding the limitations of our HEAD & TED approach, we have several points to discuss. First, the HEAD approach demonstrates the ability to distinguish anomalies that notably deviate from force field refined conformations. However, as the degree of anomaly decreases, the discriminative power drops. This reduction may stem from the energy partitioning process during the training of ANI-2x. Specifically, ANI-2x predicts the energy for each atom of a molecular conformation and combines these predictions to obtain the total energy of the molecule through simple algebraic summation. All parameters are trained to minimize the difference between the predicted total energy and the ground truth total energy. However, this process does not adequately train the energy distribution process, which poses a potential risk for our HEAD approach. Specifically, if an atom actually has high energy but that energy is distributed among its neighboring atoms, it may go undetected by the HEAD system. The risk of having undetected high-energy atoms caused by this issue is high when the total energy of the molecule is low. This is consistent with the observation in Figure 3b that the discriminative power of HEAD decreases along with the decrease of ΔE.

To address this limitation, we introduced the concept of information entropy as a metric for assessing the level of energy distribution among an atom and its spatial neighbors (details in Method section 4.1.5). By identifying conformations with high entropy and considering these conformations as invalid, we can detect previously missed abnormal conformations, as demonstrated in Figures 5a to 5d. From the perspective of weighted F1 scores (Figure 5e), we observed an improvement in HEAD’s discriminative power for conformations with small differences from force field refined ones (i.e. low ΔE). However, this improvement is achieved at the cost of reduced F1 scores for high ΔE conformations, as shown in Figure 5e. The trade-off associated with this entropy-based method highlights the importance of developing solutions from a more fundamental level, such as integrating learnable parameters into the atomic energy summation component of the ANI-2x architecture. This represents a promising direction for future improvements of HEAD approach.

**Figure 5.**
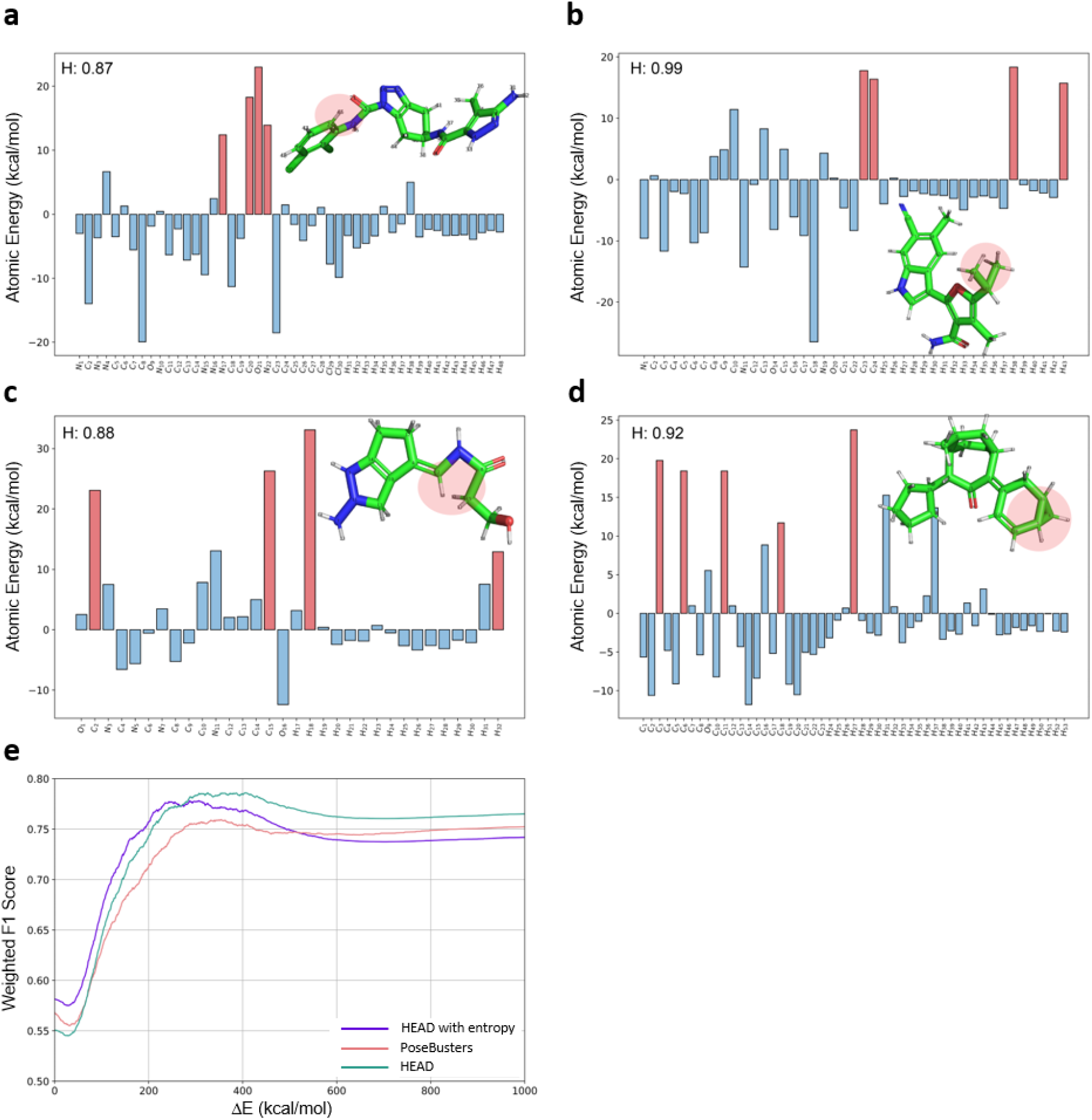
The impact of incorporating information entropy into the HEAD approach for identifying abnormal molecular conformations. (a-d) Examples of conformations identified as having high information entropy, which were previously undetected by the HEAD method. The information entropy is denoted as H. (e) Weighted F1 scores demonstrating the performance trade-off of introducing entropy-based metric. While the discriminative power of HEAD improves for conformations with low ΔE, there is a corresponding decrease in weighted F1 scores for conformations with high ΔE.

Regarding the limitations of TED, although it demonstrates the ability to distinguish molecular conformations with high strain energies (Figure 3b), its discriminative power is mild. This may be due to TED’s focus on assessing the quality of torsions without considering intramolecular interactions, which can offset the energy penalties associated with unfavorable torsion angles and thereby reduce the overall strain energy of the conformation. Extended Data Figure 4 presents an example where molecules adopt conformations with high torsion energy. However, the corresponding torsion angles in this example facilitates the formation of intramolecular hydrogen bonds, which stabilized the conformation. These considerations serve as a reminder to users of TED: when a molecular conformation is deemed irrational by TED, it indicates only that the conformation has high torsion energy. Users should further examine intramolecular interactions for a comprehensive evaluation of strain energy.

Another potential direction for improving TED is the data augmentation of the torsion energy prediction model (i.e. TED-Model). Specifically, the input conformers used during training and inference are prepared using the procedure indicated by the red arrows in Extended Data Figure 5a. This procedure involves an initial sampling to obtain no more than 20 initial conformations driven by ConfGenX^32^, followed by a systematic enumeration of torsion angles through the rotation of the target rotatable bond for each initial conformation. During model training, this enumeration is then followed by a minimum pooling operation that reduced a set of 20 × 24 conformations to a series of 24 conformations. This pooling process ensures a one-to-one correspondence between the 24 conformations and the corresponding series of 24 torsion energy values, as outlined by the green arrows in Extended Data Figure 5a. The same sampling and minimum pooling procedures are applied to a specific torsion fragment during model inference. Notably, the minimum pooling operation can be omitted during the preparation of the training data. This omission would result in multiple series of 24 conformations being linked to a single series of 24 torsion energy values. This data augmentation strategy has the potential to accelerate the inference process by eliminating the need of multiple initial conformations of the input torsion fragment. We tested this approach and observed that the data-augmented version of the torsion prediction model is five times faster than the model without data augmentation (Supplementary Information Part 4). It also achieved a Pearson correlation of 0.82 on DFT-5K, which is only slightly lower than the 0.84 correlation of the non-augmented version.

Lastly, it’s important to note that our HEAD & TED approach evaluates the quality of 3D conformations of AI-generated molecules without considering the protein pocket. Torsion energies can be influenced by the electrostatic potential or dielectric constant of the environment^33^. For instance, high torsion energy resulting from lone pair electron repulsion may be mitigated by a nearby positive charge^34^. However, no mature technology currently exists to precisely quantify this influence, posing a challenge for the accurate estimation of ligand torsion energy in the binding pocket. Since the electrostatic potential on the surface of a pocket can be calculated^35^ or observed through experimental electron density^36^, an ideal ligand torsion energy prediction model would incorporate such information. Our TED Model, trained with implicit water as a solvent, complements Rai’s vacuum-based torsion energy predictions^27^ in terms of the environmental dielectric constant, and together they provide a foundation for the future development of torsion energy prediction models sensitive to binding pockets.

## 4. Methods

Our conformation evaluation system is comprised of two key components: (1) atomic energy-based validity assessment by HEAD and (2) a torsion energy-based rationality assessment supported by TED. In this section, HEAD is described by introducing MLFF architecture and element-wise energy thresholds statistically established from a various of reference dataset. TED is described from the perspectives of training data preparation and torsion energy prediction model development.

### 4.1 HEAD Development

#### 4.1.1 Preliminary: machine learning force fields

The overview of HEAD is shown in Figure 1a. Before diving into HEAD method, we briefly introduce MLFFs as the preliminaries. More comprehensive introductions can be found in these review papers^37-39^. MLFFs aim to predict the total energy E^pred^ for a given molecule conformation {(R_i_, S_i_)}_i=1,2,‥,N_ with N atoms, where 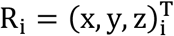 represents ith atomic positions, and S_i_ represents the scalar feature of ith atom, e.g., atomic number. The models are trained on a large scale of conformations to fit model outputs with total energies obtained from DFT calculations in a supervised manner by minimizing a loss function that typically measures the distance between the predicted energies and DFT energies also with other quantities, such as atomic forces, incorporated^19,40-43^. The nearsightedness principle is often assumed for MLFFs, where for ith atom, a receptive field of its neighboring atoms within a cutoff radius is only considered to construct ith atomic environment feature either by pre-designed symmetry-invariant functions^44-46^ or symmetry-equivariant message-passing neural networks^47,48^ with learnable features. Later, MLFFs output a set of atomic energies (also called site energies in some literature^49^) corresponding to each atomic environment feature and the total energy can be obtained by a summation over all atomic energies (Eq.1).

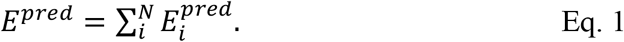

Such locality property establishes a mapping of each atomic environment feature to its corresponding atomic energy, which enables a liner scaling in system size of different molecules.

#### 4.1.2 Atomic energy extraction

HEAD is built upon MLFFs with one key difference: instead of extracting the total predicted energy E^pred^ of a given molecular conformation, it outputs the local atomic energies {E_i_^pred^}i=1,2,…,N for all atoms.

It is important to note that when projecting real space into energy space, all types of unrealistic geometries (e.g., steric clashes, uncommon molecular fragments, twisted structures, misplacement of hydrogen atoms, and excited states) can be reflected as high-energy signals in energy space compared to their realistic counterparts. This offers a comprehensive and rapid assessment, regardless of the unrealistic scenarios present in real space.

In this work, we utilize the ANI-2x model^18^, which is a variation of the prominent Behler-Parrinello neural network potentials, with modified symmetry functions. ANI-2x is designed for accurate and fast prediction of molecules composed of C, H, O, N, S, F, and Cl, covering a large set of small molecule drugs. Notably, the HEAD approach can, in principle, be applied to other MLFFs.

#### 4.1.3 Statistic element-wise energy thresholds

To distinguish high atomic energies resulting from unrealistic conformations from those of realistic conformations, we establish element-wise energy thresholds. These thresholds are derived from element-wise distributions in energy space, generated by applying MLFFs and extracting atomic energies of molecules in various molecular datasets.

Specifically, we first applied the ANI-2x model to three high-quality datasets: QMugs, OrbNet, and QM9^50-52^. The QMugs dataset contains approximately 2 million optimized molecular conformations that are biologically and pharmacologically relevant, generated using the semi-empirical GFN2-xTB method. We used 1.1 million molecules composed of C, H, O, N, S, F, and Cl from OrbNet dataset. The QM9 dataset is a widely used benchmark for MLFFs and includes 134,000 stable organic molecules made up of C, H, O, N, and F.

We then grouped the atomic energies by element type, and the element-wise distributions are presented in Supplementary Information Part 1. The energy thresholds are initialized based on the computed elbow points derived from the element-wise atomic energies. These thresholds are subsequently refined using a grid search to maximize the average weighted F1 score on the GM-1K dataset. The details of GM-1K dataset are described in Method 4.3.

#### 4.1.4 HES: high-energy score

A high-energy score (HES) is given to quantitatively measure the level of invalidity. For a given conformation, its HES is defined as,

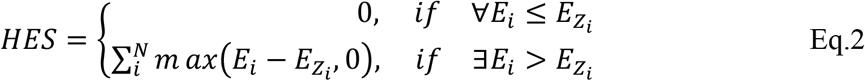

where Z_i_ is atomic number for ith atom, 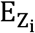 is the atomic energy threshold for that element, N is the number of atoms in the conformation. Consequently, HES is zero for valid conformation and is larger if the given conformation is more deviated from its valid counterpart. Moreover, HES can alleviate corner-case issue existed in typical threshold-based method.

#### 4.1.5 Information Entropy

We established an undirected graph for a conformer, with vertexes representing all atoms with individual atomic energy (i.e. *E*_*i*_) larger than a pre-defined value *E*_*c*_ (*E*_*c*_=10 kcal/mol in this work). Two vertexes are linked by an edge if the distance of the two atoms is less than 2 Å. For each component which contains at least two linked vertexes, its sub-region energy is calculated by summing all the atomic energies of the vertexes in it. The component with the highest sub-region energy undergoes an information entropy calculation (as described in Eq. 3 and Eq. 4), and its information entropy is then used to represent the information entropy of the entire conformer.

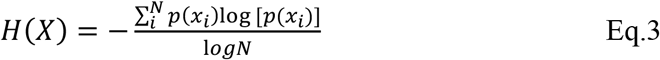

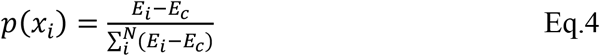

Here, H(X) represents the normalized information entropy for component X which is composed of atoms *x*_i_ satisfying *E*_*i*_>*E*_*c*_. *N* denotes the total number of atoms that satisfy this condition in component X.

#### 4.1.6 Classification Criteria for Conformation Validity

For a given conformation, it is considered as invalid if it meets any of the following two criterion. First, it contains any atom with HES larger than the atom’s element-wise energy threshold. Second, for the conformation passing the first criteria, if its information entropy is larger than 0.8, it is considered as invalid.

### 4.2 TED approach

#### 4.2.1 Torsion Energy Label Preparation

Data preparation commenced with the fragmentation of molecules. This process isolates fragments containing the rotatable bond of interest, while preserving a minimal yet suitable chemical environment around it. The fragmentation process was conducted following the methodology established by Rai (2019)^53^. It began with identifying a quartet of adjacent atoms that define the dihedral angle for the rotatable bond. This initial quartet was then expanded to include any atoms directly bonded to it. If these additional atoms belonged to predefined structural groups—such as rings or amides—the entire group was included to maintain structural integrity. For cyclic structures, substituents at the ortho position of the quartet were preserved due to their potential interactions that could influence torsion energy. Terminal carbons with unfulfilled valence electrons were capped with hydrogen atoms, while terminal heteroatoms were capped with methyl groups. The segmentation process, illustrated in Extended Data Figure 6, decomposes a molecule into a set of smaller molecules (hereafter referred to as torsion fragments), each representing a fragment of the original structure containing one rotatable bond of interest (hereafter referred to as target rotatable bond) and its local chemical environment. This method was applied to our in-house molecular database, which contains over 2 billion molecules. After decomposition, we identified more than 100 million torsion fragments, which were then sorted by their frequency of occurrence. From this sorted list, the top 100,000 torsion fragments were selected for torsion energy surface calculations of its target rotatable bond.

To characterize the torsion energy surface associated with the target rotatable bond in a torsion fragment, it is essential to identify the lowest energy among the conformers that share the same dihedral angle. This ensures that the surface accurately reflects the influence of torsion angles on energy, without being affected by high-energy conformations of other groups in the torsion fragment. This objective was achieved through a three-step sampling approach, as illustrated by the green arrows in Extended Data Figure 5a. Initially, no more than 20 initial conformations were randomly generated for each torsion fragment using ConfGenX, providing a diverse set of initial conformations to minimize the risk of trapping in local minima during subsequent steps. The target rotatable bond in each initial conformation was then repeatedly rotated in increments of 15 degrees, resulting in 24 conformations per initial conformation. However, this procedure treats the moving parts of the molecule as rigid, which may lead to steric clashes. To address this issue, the conformations underwent MMFF94 force field optimization with the target torsion angle fixed. Subsequently, to ensure that high torsion energy was influenced solely by the improper torsion angle and not by the high-energy conformations of other groups within the same torsion fragment, molecular dynamics simulations were conducted with the target torsion angle held constant. These simulations, powered by GFN-FF^54^ at 400K and conducted using xTB^55^ program, generated up to 100 conformers per input. As a result, we accumulated no more than 48,000 conformations per torsion fragment (i.e., 20 x 24 x 100). The conformers were subsequently grouped into 24 categories based on the dihedral angle values of the target rotatable bond. For each group, GFN1-xTB optimizations were performed, and the optimized conformations with the lowest GFN2-xTB single-point energy was selected as the representative conformation for that dihedral angle. Further refinement of these representative conformers was conducted using B3LYP-D3(BJ)/def2-SVP optimizations using ORCA^56^, with the dihedral angles maintained as fixed. This was followed by single-point energy calculations performed at the revDSD-PBEP86-D3(BJ)/def2-TZVPP level. All calculations utilized the SMD water solvent model. The resulting double hybrid DFT level single-point energies were used to plot the torsion energy surface for the rotatable bond.

In addition to the aforementioned procedure of torsion energy surface calculation, we also developed a streamlined version to produce large amount of data for model pretrain. It retained the key elements of the original approach but excluded the molecular dynamics and DFT components, providing GFN2-xTB level torsion energy surfaces. We applied the streamlined method for 6 million molecules of our 100 million torsion fragments library. These GFN2-xTB level data were used as complementary of the DFT-level data during model training.

We have presented two approaches for calculating the torsion energy of a target rotatable bond in a torsion fragment. From a model training perspective, both approaches serve to generate labels.

#### 4.2.2 Torsion Fragment Feature Extraction

For torsion fragment conformational feature extraction, our methodology involves characterizing the target atom quartet and the local chemical environment surrounding the target rotatable bond.

We used five key features to characterize a quartet of atoms. For an atom quartet represented as *abcd*, where *a, b, c*, and *d* denote individual atoms and *bc* denotes the rotatable bond of interest, the features are defined as follows: (1) the dihedral angle between plane *abc* and *bcd*; (2) the distance between atoms *a* and *d*; (3) the distance between atoms *b* and *c*; (4) the product of the atomic numbers of atoms *a* and *d*; (5) the product of the atomic numbers of atoms *b* and *c*. This results in a 5-dimension feature for atom quartets.

To extract features of the local chemical environment of a rotatable bond, we adapted atomic environment vectors (AEVs)^19,43^, which are designed to encode the local chemical environment of a target atom. Given our focus on rotatable bonds, we modified the AEV approach by incorporating distance, angular, and dihedral symmetry functions (SFs) to encode the local chemical environment of rotatable bonds.

Distance symmetry function encodes the distance between the center of bond with any other atoms of the conformer. The formula for distance feature extraction is Eq.5.

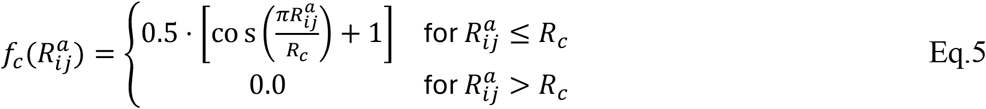

Here, we denote the center of the rotatable bond of interest as point *i* and any arbitrary atom in the conformer as point *j*. R_c_ is a threshold that takes values from the set {1.5, 2.0, 2.5, 3.0, 4.0, 6.0, 10.0}. *R*_*ij*_^*α*^ denotes the distance between atoms *i* and *j*, with the superscript *α* indicating the atom types: H, C, N, O, F, S, Cl, along with two formal charges, resulting in a total of nine element types. The distance symmetry function 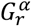 for each type is computed using Eq.6.

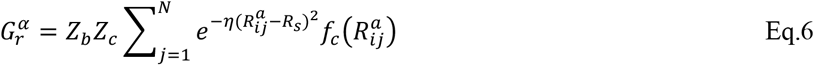

Here, *Z*_*b*_ and *Z*_*c*_ represent the atomic numbers of the atoms at both ends of the bond of interest, while *η* and *R*_*s*_ are set to 10^-4^ and 0, respectively. This results in a 63-dimension distance feature (9 element types × 7 *R*_*c*_ values).

The angular symmetry function (Eq.7) encodes the angles formed by the center of the rotatable bond and any two other atoms in the conformer.

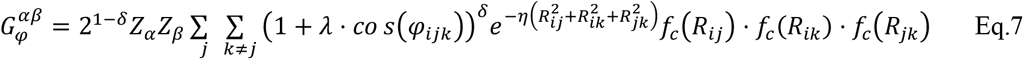

Here, *Z*_*α*_ and *Z*_*β*_ represent the atomic numbers of the atoms at both ends of the bond of interest. The subscript *i* represents the center point of the bond in the dihedral angle, while *j* and *k* are any two atoms in the conformation. *R*_*ik*_, *R*_*ij*_, *R*_*jk*_ represent the distances between subscript specified atoms, and *ϕ*_*ijk*_ denotes the angle formed by the edges *R*_*ij*_ and *R*_*ik*_. The parameters *δ, λ, η* are set to 0.5, 0.5, and 10^-4^, respectively, with *R*_*c*_ valued at 4.0. Given nine types of elements, there are a total of 45 combinations of atom pairs (CC, CN, CO, …, CS, CH), resulting in a 45-dimension angle feature (1 *R*_*c*_ value × 45 combinations of atom pairs = 45).

The dihedral symmetry function (Eq.8) encodes the torsion angles of the rotatable bond of interest involving any two other atoms in the conformer.

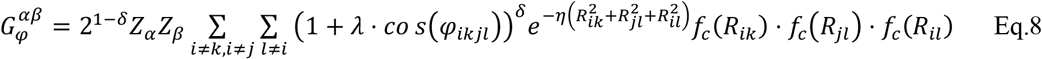

Here, we denote atoms *k* and *j* as the two terminal atoms of the bond on interest. For any other two atoms, *i* and *l*, atoms *i, k, j, and l* form quadruplets, with *R*_*ik*_, *R*_*jl*_, *R*_*il*_ represent the distances between subscript specified atoms, and *ϕ* _*kjl*_ representing the dihedral angle of planes *ikj* and *kjl. R*_*c*_ takes values from the set {2.5, 3.5, 5.0, 10.0}. The parameters *δ, λ, η* are set to 0.5, 0.5, and 10^-4^, respectively. Given nine types of elements, there are a total of 45 combinations of atom pairs. This results in a 180-dimension dihedral angle feature (4 *R*_*c*_ values × 45 combinations of atom pairs = 180).

The hyperparameters *R*_*c*_, *δ, λ*, and *η* were configured in accordance with the parameters established in previous AEV studies^19,43^.

The 63-dimension distance, 45-dimension angle, and 180-dimension dihedral angle features for rotatable bonds and 5-dimension feature for atom quartets are concatenated to yield a 293-dimension feature which is used as input for the model.

#### 4.2.3 Torsion Energy Prediction Model Training

Our torsion energy prediction model (i.e. TED-Model) is designed to predict energy values for an input series of 24 conformations with target torsion angles at intervals of 15 degrees. The input conformations should be obtained using inexpensive methods, such as MMFF94 force field, while the output torsion energy must exhibit a high level of accuracy, ideally at DFT level. Otherwise, the model would be unnecessary, as an inexpensive torsion energy surface can be generated simultaneously with the generation of input conformations. The schematic flow indicated with red and green arrows in Extended Data Figure 5a summarizes the feature extraction and torsion energy labeling process, respectively. This process results in a series of 24 conformations with torsion angle at 15° intervals and a series of 24 energy values as labels.

The data supporting model training consists of two datasets. The first dataset is a double hybrid DFT-level torsion energy dataset involving 100,000 torsion fragments, while the second dataset is a GFN2-xTB level torsion energy dataset involving 6 million torsion fragments.

The training process was divided into two phases: pretraining and fine-tuning. The complete set of GFN2-xTB level data, along with 10% of the double hybrid DFT data, was utilized for pretraining, while the remaining 90% of the double hybrid DFT data was reserved for the fine-tuning phase. In each phase, the data was split into training and test sets using a 8:2 ratio. To prevent information leakage, no molecules in the test set shared more than 0.5 Tanimoto similarity with any molecules in the training set, based on ECFP4 fingerprints^57^.

The input for the model consists of a series of 24 conformations, with each conformation represented as a 293-dimension vector. This represents the input as a two-dimensional matrix of size 24 × 293. To effectively capture the relationships among the input conformations, we employ a self-attention framework. The output of self-attention module is then flattened and input into a linear-layer module which generate predictions for each conformation, resulting in a final output of size 24 × 1. The model architecture is shown in Extended Data Figure 5b.

The input matrix M is of size 24 × 293, where each row represents a conformation, and the rows are sequenced in ascending order according to dihedral angle values of specified atom quartet. M undergoes an attention^58^ transformation using Eq. 9.

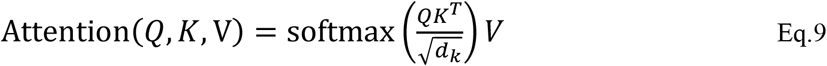

This process generates a new matrix representation of size 24 × 293 that encapsulates the internal relationships among the 24 conformations. Thereafter, we apply a flatten operation to compress this matrix into a 7032-dimension vector. Then the 7032-dimension vector is put into linear layers module which consists of seven layers with number of neurons arranged as follows: 3516, 1758, 879, 293, 146, 73, and 24. Each linear layer uses ReLU as active function and employs batch normalization strategy, with a dropout rate of 10% applied during training. The output of the linear layers is a 24-dimension vector. The model’s loss function is defined as the Mean Absolute Error (MAE) of the energy values, expressed as Eq. 10.

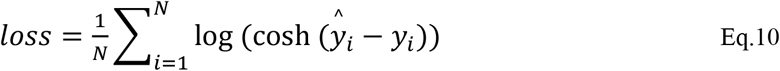

Here, 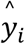 and *y*_*i*_denote the predicted energy values and energy value label for conformer *i*, respectively.

Adam was used as optimizer with initial learning rate set as 0.001. Early stop strategy was used to avoid overfitting.

#### 4.2.4 Classification Criteria for Conformation Rationality

This process is illustrated in Extended Data Figure 5c. For a given conformation, it is first decomposed into several torsion fragments using the method described in Section 4.2.1. Subsequently, for each torsion fragment, a series of 24 torsion energy values corresponding to 24 torsion angles at 15°intervals is p redicted. The torsion energy surface is then generated by smoothing the 24 discrete points using linear interpolation in SciPy package^59^. This method ensures that the interpolated values remain within a reasonable range. It helps avoid the extreme values or oscillations that can arise from cubic interpolation, which may lead to unrealistic dips below zero or spikes in energy. The torsion energy for each fragment is determined by checking the corresponding energy value for the torsion angle adopted in that fragment against the torsion energy surface. If any of the torsion fragments derived from the given conformation has a torsion energy exceeding 2 kcal/mol, the conformation is classified as irrational. This threshold of 2 kcal/mol is based on findings reported in a previous study^53^.

### 4.3 Datasets

#### 4.3.1 GM-5K

The GM-5K dataset contains 5,000 molecules randomly selected from AI-generated molecules produced by five models: Lingo3DMolv2, Pocket2Mol, PocketFlow, TargetDiff, and PMDM. Each model was set to generate 1,000 molecules for each of the 102 targets in the DUD-E dataset. We randomly select 1,000 molecules from each model, resulting in the final GM-5K dataset of 5,000 molecules.

For each conformation in the GM-5K dataset, geometry optimization using the MMFF94 force field was performed. This is followed by QM-level single-point energy calculations (revDSD-PBEP86-D3(BJ)/def2-TZVPP) for both the pre-optimized and post-optimized conformations. The energy differences, calculated as ΔE = E_ori_^DFT^ - E_opt_^DFT^, serve as indicators of conformation quality.

#### 4.3.2 GM-1K

The GM-1K dataset is created using the same methodology as GM-5K, but it includes only 1,000 molecules that do not share similar counterparts with GM-5K. Similarity is determined using Tanimoto similarity for ECFP4 fingerprints; molecules are considered similar if they have a Tanimoto similarity greater than 0.5.

#### 4.3.3 DFT-5K

The approach described in Method section 4.2.1 and illustrated in Extended Data Figure 6 was initially applied to our in-house molecular database, which contains over 2 billion molecules, resulting in the identification of more than 100 million torsion fragments. These fragments were then sorted by their frequency of occurrence. From this sorted list, the most frequently occurred 100,000 torsion fragments were selected for torsion energy surface calculations at the DFT level and used for TED-Model training. The DFT-5K dataset consists of 5,000 unique torsion fragments selected from the previously mentioned 100-million torsion fragment library, ensuring that no DFT-5K torsion fragment has its Bemis–Murcko scaffolds appearing in the DFT level training set for TED-Model.

## Data and Code Availability

The GM-5K, GM-1K, and DFT-5K datasets are available via the https://github.com/stonewiseAIDrugDesign/HEAD_TED.

We obtained the models for Pocket2Mol, TargetDiff, PocketFlow, and PMDM from their official GitHub repositories and used them for molecule generation. For Lingo3DMolv2, we utilized the online service accessible at https://sw3dmg.stonewise.cn to generate molecules. Our source code for the HEAD and TED models is publicly available on our GitHub repository at https://github.com/stonewiseAIDrugDesign/HEAD_TED.

## Acknowledgements

This study was funded by the National Key R&D Program of China (grant no. 2022YFF1203004 received by B.H.). This work was also supported by the Beijing Municipal Science and Technology Commission (grant no. Z241100007724005 received by B.H.).

## Author contributions

B.H., F.Z, and W.Z conceived the study. B.X developed HEAD. F.F. and X.M developed TED. W.Z provided instructions for artificial intelligence modelling. F.Z. provided instructions on QM calculations. H.Z. provided instructions for torsion fragment construction. F.F., Q.X., H.W., W.F., J.D. supported the evaluation of AI models. Z.L. and B.X. supported forced field-based optimization.

---

**Extended Data Table 1.** Speed comparison for HEAD and PoseBusters on LigBoundConf, CSD and GM-5K benchmark dataset. HEAD was run on single NVDIA GeForce RTX 3090 GPU, while PoseBusters was run on single CPU as its current version only supports CPU.

|  | # conformations | Time Cost (min) |  |
| --- | --- | --- | --- |
|  |  | HEAD | PoseBusters |
| LigBoundConf | 6209 | 1.7 | 49 |
| CSD | 159948 | 47 | 2353 |
| GM-5K | 5000 | 1.6 | 40 |

**Extended Data Figure 1.**
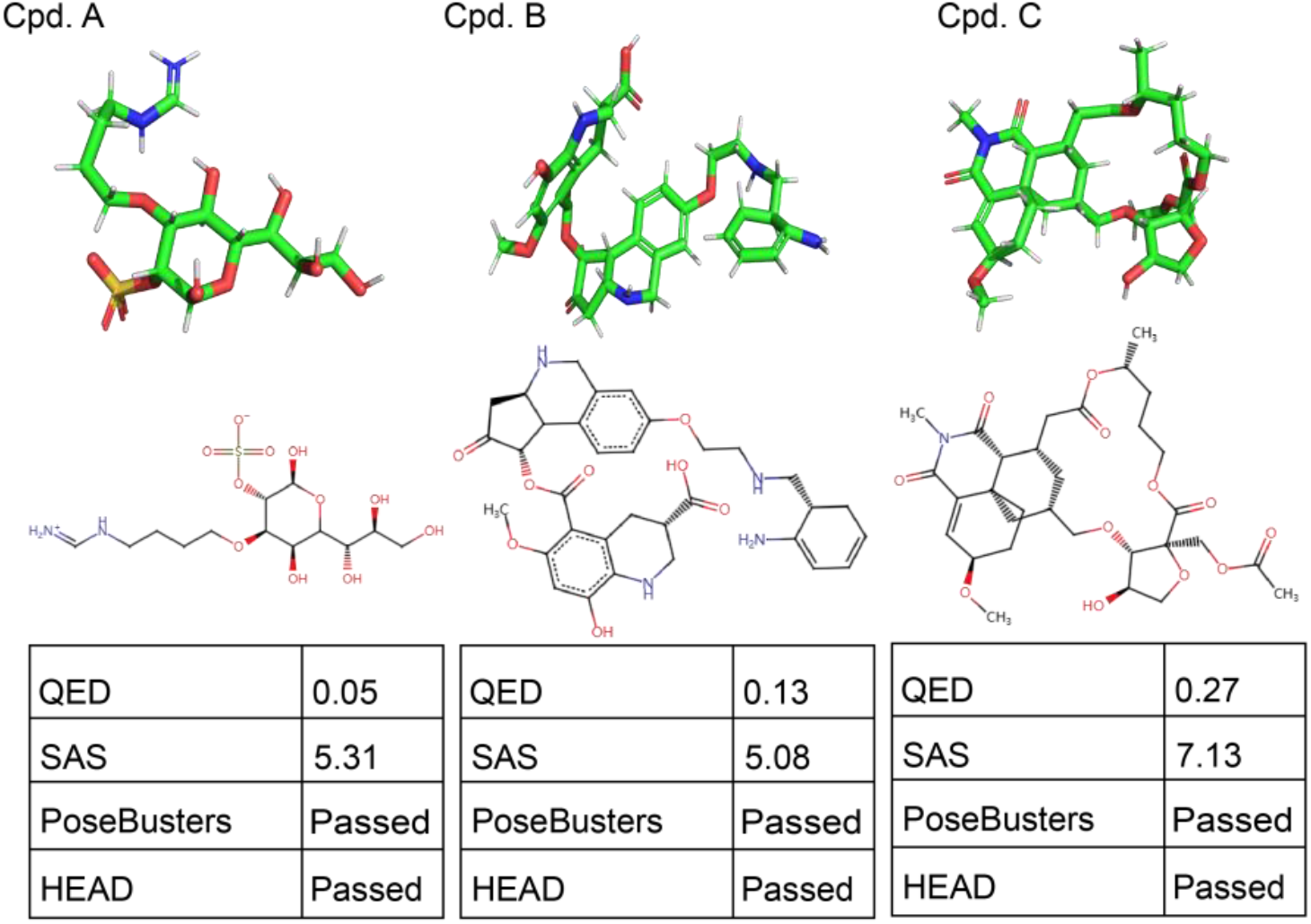
AI generated molecules that passed the PoseBusters and HEAD tests but are not suitable as drug candidates. Three specific compounds—Cpd. A, Cpd. B, and Cpd. C—are displayed. Although these compounds successfully passed the PoseBuster and HEAD tests, they exhibit unfavorable QED or SAS. Including these molecules may create a misleading impression of high conformational quality; therefore, they should be excluded from assessments of conformational quality.

**Extended Data Figure 2.**
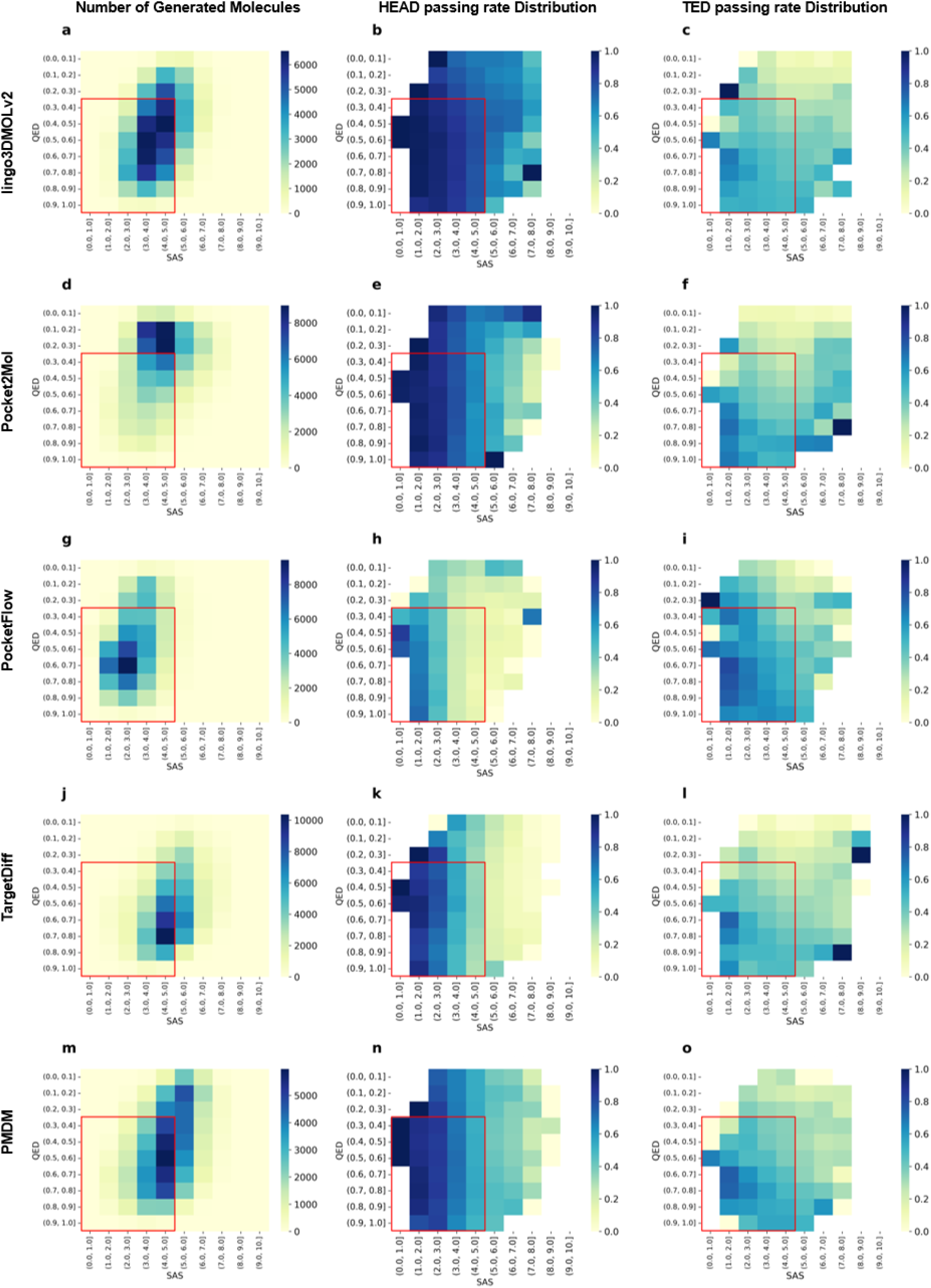
Distribution of HEAD and TED passing rates against QED and SAS for molecules produced by various models. Panels a-c, d-f, g-i, j-l, m-o indicates results for Lingo3DMolv2, Pocket2Mol, PocketFlow, TargetDiff, and PMDM, respectively. The drug-like region, defined with QED score higher than 0.3 and SAS less than 5, is squared with redlines. For each model, number of generated molecules, distributions of HEAD and TED passing rates are shown as heatmap.

**Extended Data Figure 3.**
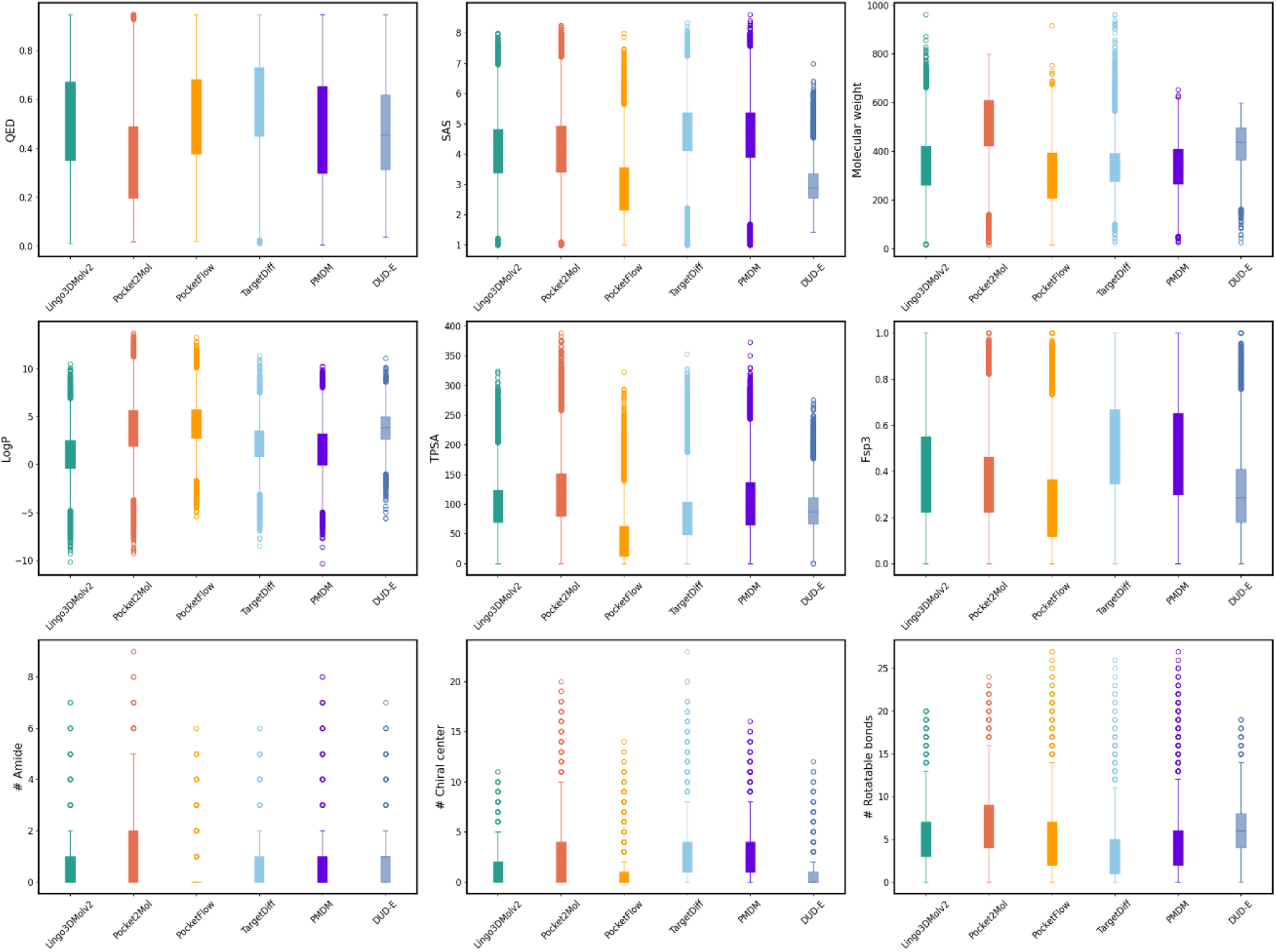
Property distributions of AI generated molecules for DUD-E targets. Active molecules in DUD-E are displayed as references. Properties including QED, SAS, molecular weight, LogP, Topological polar surface area (TPSA), fraction of *sp*^3^ carbons (Fsp3), number of amide groups, number of chiral centers, and number of rotatable bonds are analyzed.

**Extended Data Figure 4.**
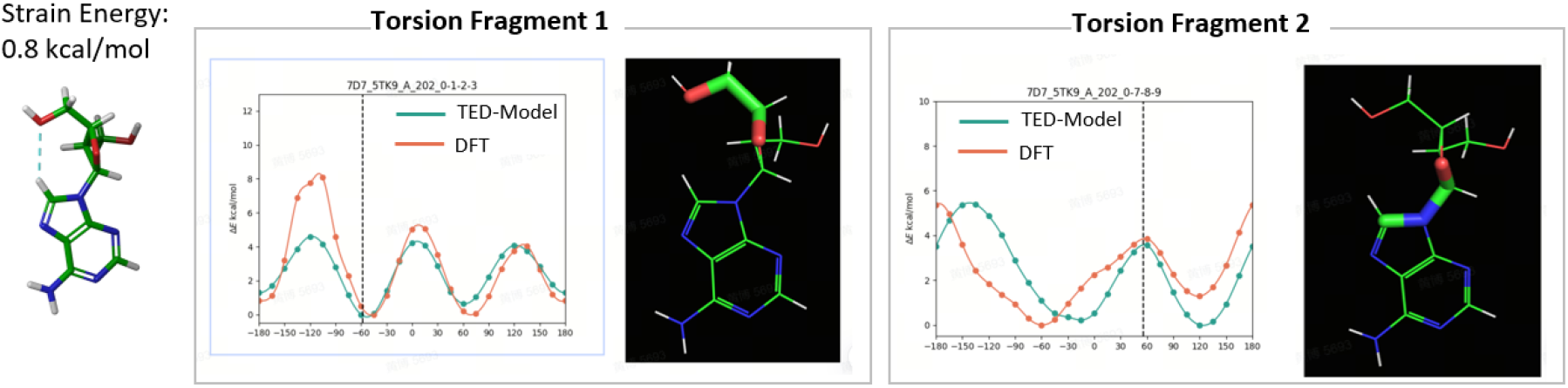
Intramolecular interactions stabilizing unfavorable torsion angles. Strain energy was sourced from the LigBoundConf database. An intramolecular hydrogen bond stabilizing torsion fragment 1, which adopts an unfavorable angle, is illustrated using light blue dashed line. The torsion energy surfaces for each torsion fragment are shown, with torsion angle values adopted in the conformation marked by black dashed lines. Atom quartets are represented as sticks.

**Extended Data Figure 5.**
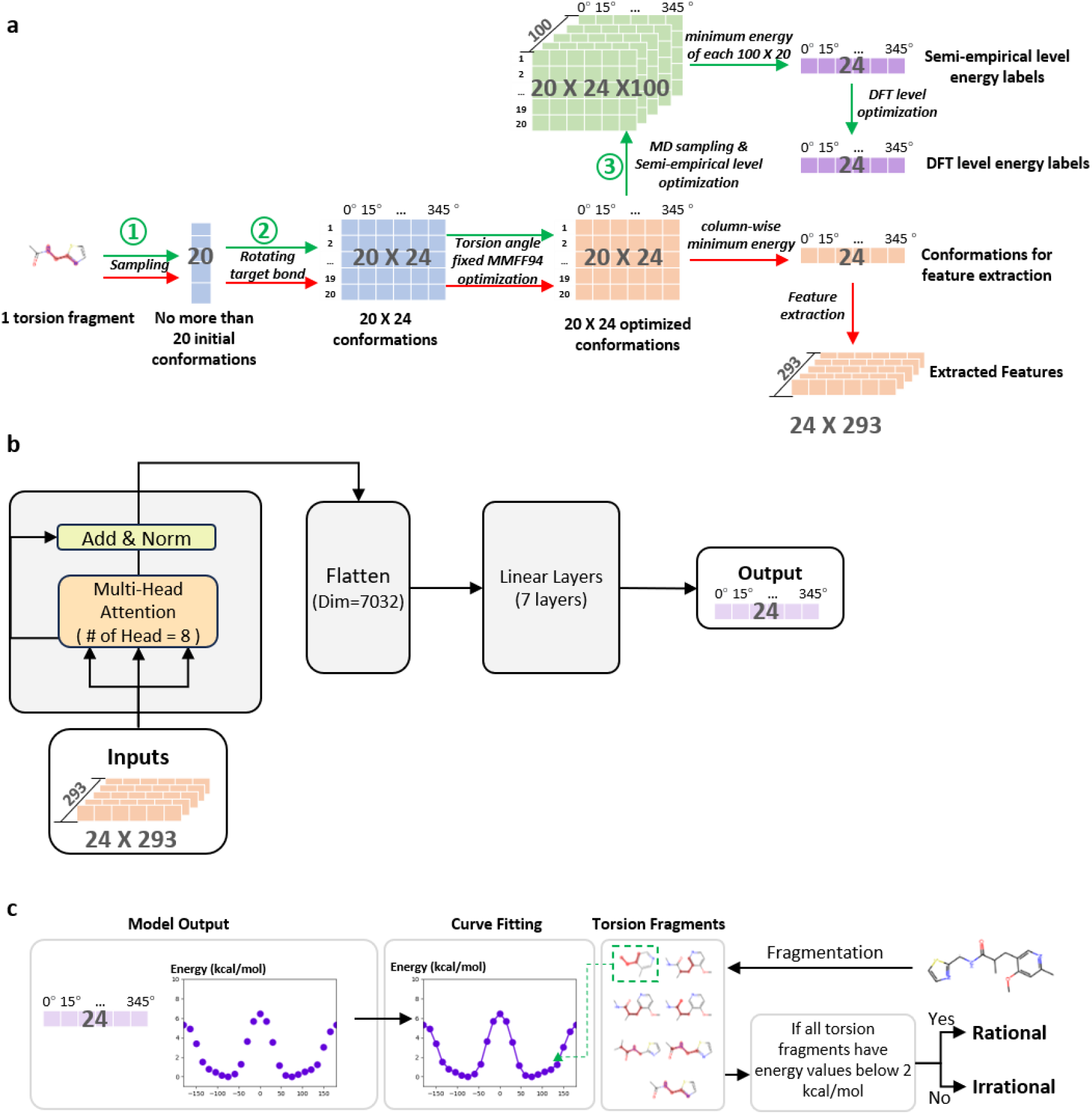
Development of TED. (a) Schematic diagram illustrating the generation of features and labels, indicated by red and green arrows, respectively. Energy label generation consists of three rounds of sampling, each represented by an index within a green circle: 1. initial sampling; 2. torsion angle enumeration; 3. MD sampling. (b) Architecture of the torsion energy prediction model. The linear module is comprised of seven layers with neurons organized as follows: 3516, 1758, 879, 293, 146, 73, and 24. (c) Procedure for evaluating the rationality of conformations by assessing torsion energy for each torsion fragments. Each conformation is decomposed into torsion fragments, which are then analyzed for torsion energy across 24 angles at 15° intervals. A torsion energy surface is generated by smoothing the energy values. Conformations with any torsion fragment energy exceeding 2 kcal/mol are classified as irrational.

**Extended Data Figure 6.**
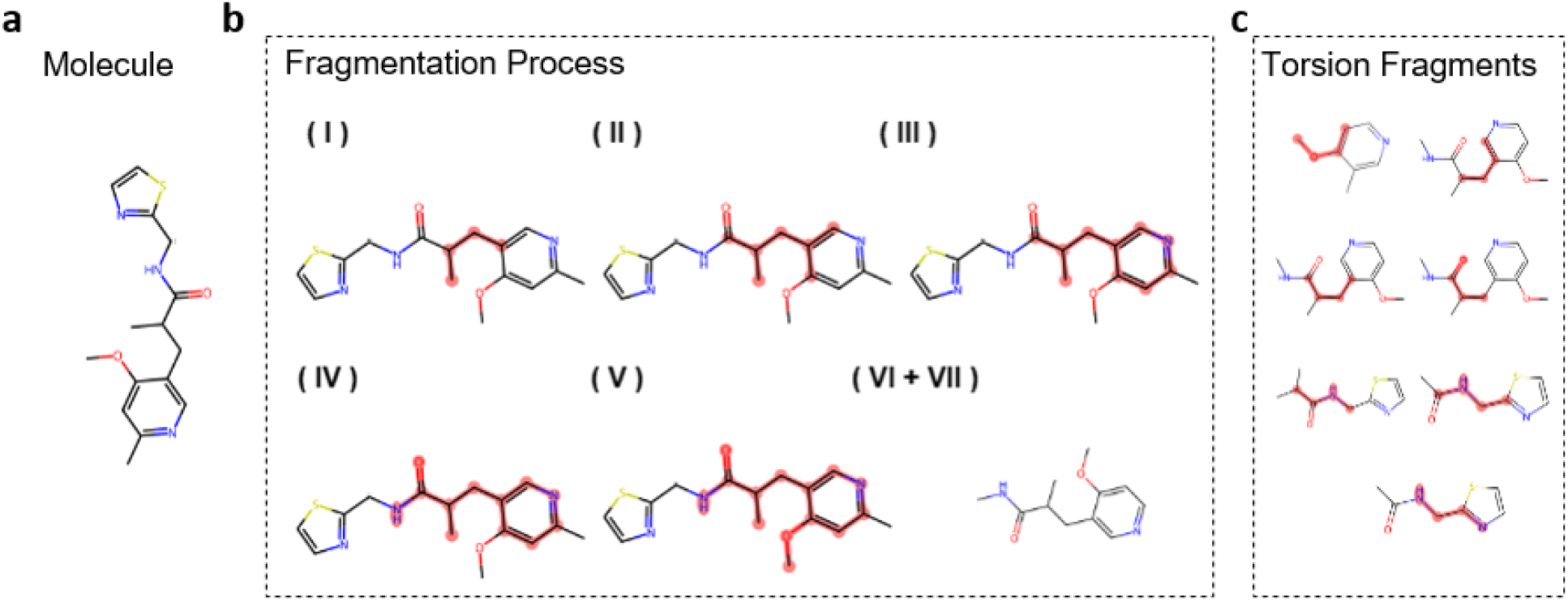
Illustration of fragmentation process for torsion fragment identification. (a) Structure of a sample molecule. (b) Step-by-step illustration of identifying one torsion fragment in the sample molecule. The fragmentation process contains seven steps. (I) Identification of the atom quartet. (II) Expansion of the atom quartet to include any atoms connected by a single bond. (III) Inclusion of atoms contributing to ring structures. (IV) Inclusion of atoms belonging amide groups. (V) Inclusion of substituents at the ortho position. (VI) Terminal capping, where heteroatoms with unoccupied valence electrons from bond cleavage are capped by methyl groups, and other atoms like carbons are terminated with hydrogens. (VII) Chirality retention, where carbon atoms that would lose chirality due to bond cleavage are terminated with methyl groups to retain their stereochemistry. (c) List of torsion fragments identified in the sample molecule.

